# Cellular Reprogramming by Notch Inhibition Regenerates Hair Cells and Afferent Neurons to Restore the Sense of Balance

**DOI:** 10.1101/2024.10.02.616053

**Authors:** Hanae Lahlou, Hong Zhu, Wu Zhou, Albert S.B. Edge

## Abstract

Sensory hair cell loss in the vestibular organs of the inner ear causes balance disorders which are essentially irreversible due to the lack of hair cell regeneration. Here, we administered a γ-secretase inhibitor to an adult mouse model of vestibular hair cell loss. The treatment regenerated type I and type II hair cells and restored canal and otolith afferent innervation, resulting in a complete recovery of rotational and translational vestibulo-ocular reflexes across all frequencies. Genetic deletion of *Notch1* in supporting cells identified Notch1 as the target of the drug. The results demonstrate that a single injection of a γ-secretase inhibitor is a viable therapy for functional restoration of the vestibular system in patients with balance disorders.

## Introduction

Loss of mechanoreceptor hair cells in the vestibular organs of the inner ear causes balance disorders that affect millions of people worldwide resulting in falls and reduced quality of life. Vestibular dysfunction is a major public health issue in the aging population, and treatments that can restore balance are lacking (*1, 2*). Hair cells are vulnerable to insults such as ototoxic agents, as well as mutations and damage due to aging (*3–6*). These insults cause irreversible damage, because of the inability of the inner ear organs to regenerate physiologically active hair cells. Vestibular hair cells reside in the semicircular canals and otolith organs of the inner ear that sense angular and linear acceleration, respectively. Two types of vestibular sensory cells, type I and type II hair cells, are interdigitated by supporting cells in a sensory epithelium (*7*) and innervated by afferent neurons (*6, 8–10*) that encode head motion signals for transmission to the central nervous system. The 2 hair cell types are innervated differently, with nerve endings comprising bouton, calyx, or both, and with regular (type II) or irregular (type I) firing patterns.

Unlike non-mammalian vertebrates that regenerate vestibular hair cells, mammals display a limited capacity for spontaneous regeneration, particularly after the early postnatal period (*11–24*). In newborn mammals, supporting cells appear to act as progenitors to regenerate type I and type II hair cells (*18, 24, 25*), which are thought to result from independent developmental trajectories from a common progenitor cell (*26*). The limited regeneration seen in the mature utricle (*15, 27*) leads exclusively to hair cells of type II phenotype. The failure to regenerate type I hair cells, along with the insufficient recovery of type II hair cells, impedes functional restoration, as both are vital for sensing head movement (*19, 24, 28–30*).

Despite extensive efforts to develop interventions that could restore the sense of balance, most approaches have not regenerated functional hair cells. Some approaches have relied on the overexpression of bHLH transcription factor, *Atoh1*, to reprogram supporting cells to hair cells in mature vestibular organs but have not restored a high level of function (*19, 31, 32*). This challenge is significant because in addition to the hair cells their innervation needs to be restored for the sensory information to be transmitted to the brain.

Here, we chose to target γ-secretase, an intramembrane protease that recognizes Notch as one of its substrates, to stimulate hair cell regeneration from vestibular supporting cells. We reasoned that inhibiting Notch could be effective (i) because Notch plays a central role in hair cell development by determining hair cell vs supporting cell fate from early sensory progenitor cells (*33–35*) and (ii) because its inhibition can regenerate hair cells in the newborn cochlea (*36–39*). We show significant regeneration of type I and II hair cells, resulting in a full recovery of vestibular function across stimulus frequencies in an adult mouse model of vestibular hair cell ablation. The new hair cells are structurally integrated into the sensory epithelium, and the treatment rescues canal and otolith function, restoring head rotation- and translation-evoked vestibulo-ocular reflexes (VORs), indicating that γ-secretase inhibitor treatment is a potential therapeutic approach to this important and widespread disability.

## Results

### Hair cell regeneration induced by γ-secretase inhibitor, LY411575, in *Pou4f3^DTR^*-ablated utricle and canal cristae

We tested inner ear injection of γ-secretase inhibitor LY411575 as a treatment for vestibular hair cell loss in adult mice. These drugs are effective tools for conversion of inner ear supporting cells to hair cells in organoids (*39*) and newborn cochlear and vestibular organs (*36, 38, 40*) but show limited efficacy in the adult cochlea (*41–43*). We chose to test the effect of the γ-secretase inhibitor in the adult vestibular system because it retains some regenerative capacity into adulthood (*44*).

We administered LY411575 by unilateral injection onto the round window membrane of DT-ablated *Pou4f3^DTR/+^* mice (*15*), 7 days after hair cell ablation (Fig. 1A). In these mice, the *Pou4f3* promoter drives expression of the human diphtheria toxin receptor (DTR) in hair cells, and administration of DT results in near-complete ablation of hair cells. We assessed regeneration in the canal and otolith vestibular epithelia by immunostaining for SRY-box transcription factor 2 (Sox2), myosin7a (Myo7a) and tenascin (Tnc) to identify supporting cells (Myo7a-Sox2+), type I (Myo7a+Tnc+) and type II (Myo7a+Sox2+) hair cells (Fig. 1B).

**Fig. 1.**
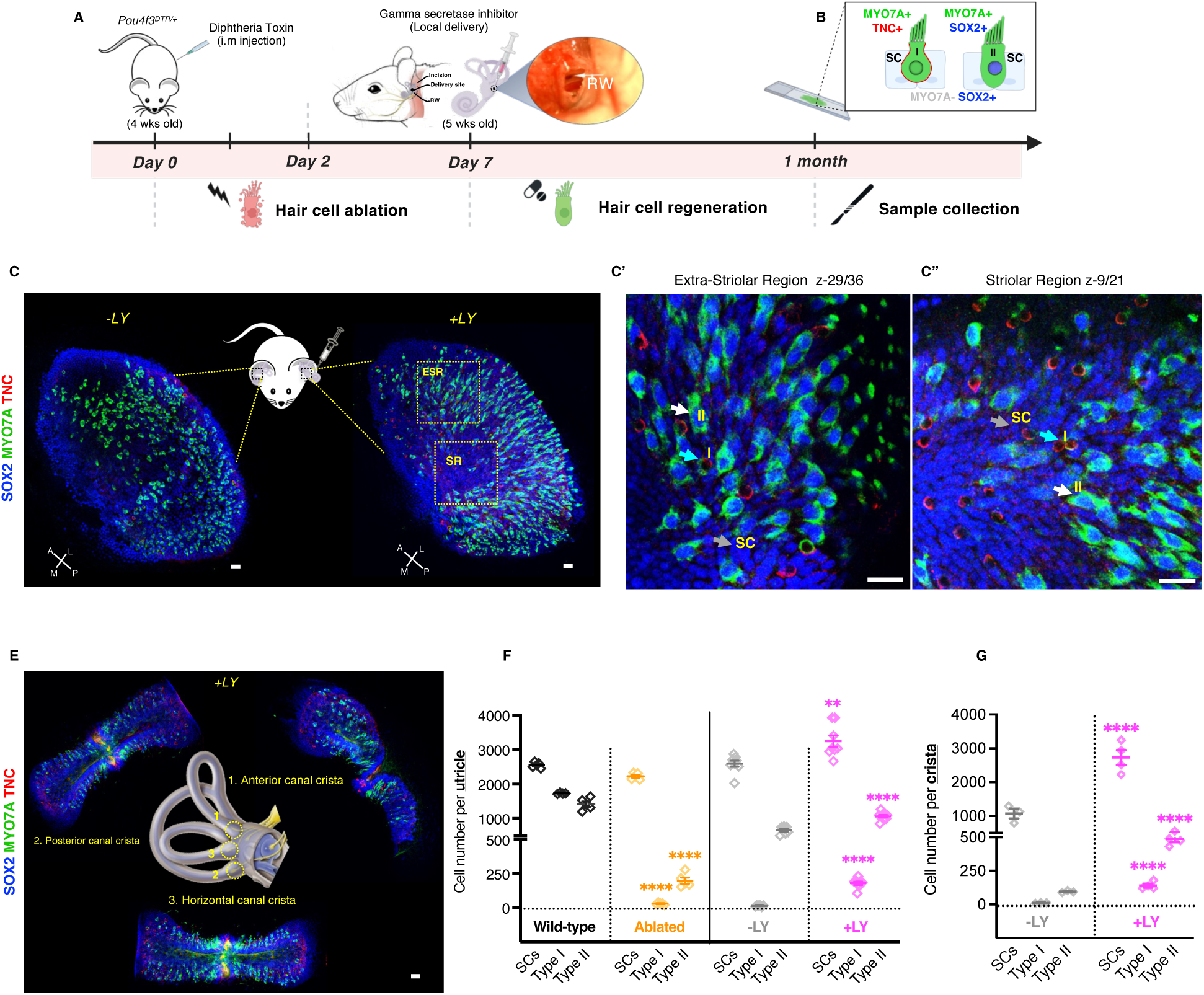
LY treatment promotes type I and type II hair cell regeneration in the vestibular organs. **(A)** Schematic of the *in vivo* approach for drug delivery in DT-pretreated *Pou4f3^DTR/+^*mice. LY was injected in the left ear via the round window (RW) 7 days after DT ablation; the contralateral ear was used as a control for spontaneous regeneration. Mice were examined 1 month after drug treatment. **(B)** Schematized transverse section illustrating type I (I, Myo7a+Tnc+) and type II (II, Myo7a+Sox2+) vestibular hair cells and supporting cells (SC, Myo7a-Sox2+). **(C-C”)** Sox2 (blue), Myo7a (green) and Tnc (red) immunolabelled utricles from untreated (-LY) and treated (+LY) ears with high-magnification images from the extrastriolar **(C’)** and striolar regions **(C”)** indicating the presence of supporting cells (gray arrows), type I hair cells (cyan arrows) and type II hair cells (white arrows). **(E)** Immunolabeled cristae from drug treated ear. **(F** and **G)** Quantification of supporting cells, type I and type II hair cells from utricles and cristae **(G)** untreated and treated ears. All data represent the mean ± SEM. \*\*\*\**p* < 0.0001 and \*\**p* < 0.01 by 1-way ANOVA with Tukey’s multiple comparisons test. Scale bars = 50-100 µm.

Examination of the utricle (otolith organ), 1 month after LY411575 treatment, (Fig. 1C) revealed a stikingly higher regeneration of hair cells than the untreated utricles (Fig. 1F) attributed to spontaneous regeneration. Moreover, while regeneration in the adult mouse utricle was limited to type II hair cells as expected (*12, 14, 17*), regeneration after LY411575 treatment comprised type I and type II cells. These cells were apparent in both striolar and extrastriolar regions of the utricle (Fig. 1C’ *and* C”).

Analysis of the anterior, posterior, and horizontal semicircular canals showed abundant Myo7a+ cells in drug-treated (+LY) ears after 1 month (Fig. 1C) whereas untreated control ears (-LY) had minimal spontaneous regeneration (Fig. 1G). Supporting cell numbers were also elevated in LY411575-treated ears in both utricle and cristae compared to controls indicating increased supporting cell proliferation following γ-secretase inhibition (Fig. 1F and G).

### Rescue of canal and otolith function by γ-secretase inhibition

To evaluate the impact of LY411575 on vestibular function following hair cell ablation, we measured vestibulo-ocular reflexes (VORs) in wild-type and DT-ablated mice with and without treatment (Fig. 2A). The VOR stabilizes gaze by producing compensatory eye movements during head motion. Canal function was evaluated using the rotational VOR (RVOR) and otolith function using translational VOR (TVOR), which respond to angular and linear acceleration respectively (Fig. 2B).

**Fig. 2.**
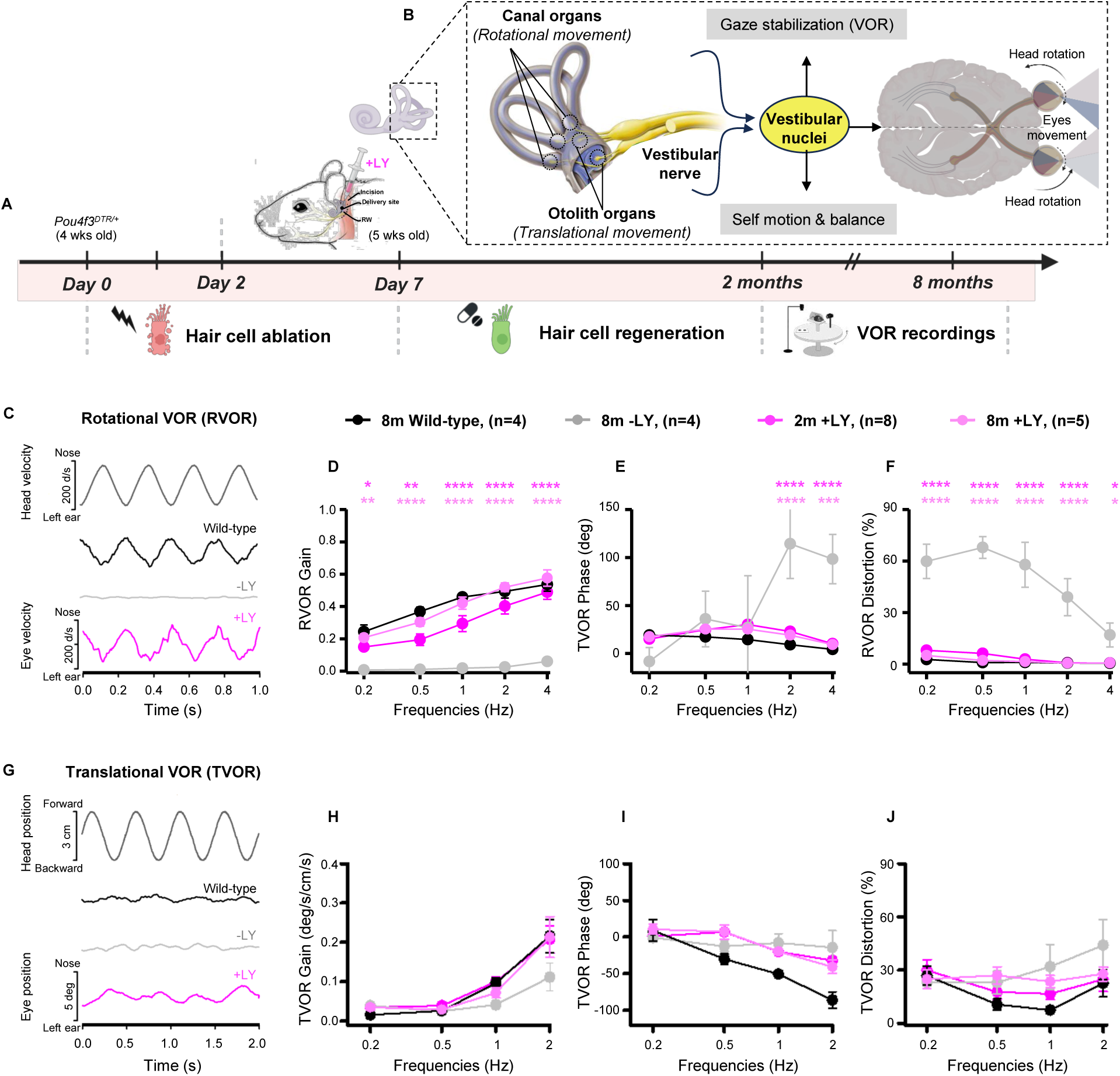
Rescue of rotational and translational VOR responses following a single dose of LY treatment. **(A)** Schematic of the *in vivo* approach for drug delivery in DT-pretreated *Pou4f3^DTR/+^*mice. Rotational and translational VORs were recorded from 2- and 8-month-old wild-type, *Pou4f3^DTR/+^* DT-ablated without drug (-LY) and *Pou4f3^DTR/+^*DT-ablated with drug treatment (+LY). **(B)** Schematic representation of vestibular signal transmission and its integration with the visual system. Vestibular inputs from the five sensory organs are transferred to the vestibular nuclei via the vestibular nerve. The vestibular nuclei then project to various brain regions involved in stabilizing the visual axis through the vestibulo-ocular reflex (VOR). **(C)** Rotational vestibulo-ocular reflexes (RVORs) to sinusoidal head rotations with representative eye velocity responses to 4 Hz head rotation at 2 months +/-LY, from wild-type (black), -LY (gray) and +LY (pink). RVOR gains **(D)** phases **(E)** and distortions **(F). (G)** Translational vestibulo-ocular reflexes (TVORs) in response to sinusoidal head translation at 2 Hz 2 months after ablation, from the same groups. TVOR gains **(H)** phases **(I)** and distortions **(J)**. All data represent the mean ± SEM. *\*\*\*\*p* < 0.0001, *\*\*\*p* < 0.001, \*\**p* < 0.01and \**p* < 0.05 by 2-way ANOVA with Tukey’s multiple comparisons; p-values reported in the graphics are from the comparisons of LY treatments to the spontaneous regeneration (-LY). See also Fig. S1 and Table S1.

Two months after treatment, LY411575-treated mice (pink lines) exhibited significant restoration of RVOR gain across all tested frequencies (Fig. 2C and D). Phase responses approached wild-type values with minimal signal distortion (Fig. 2E and F), indicating recovery of canal function. Similarly, TVOR gain responses were comparable to wild-type (Fig. 2G and H), with improved phase synchronization and low distortion (Fig. 2I and J).

At 8 months, VOR responses revealed further functional stabilization; both RVOR and TVOR gains at this time point had become indistinguishable from wild-type levels at all tested frequencies (Table S1), with normalized phases and minimal distortion. In contrast, untreated DT-ablated mice (gray lines) exhibited undetectable RVOR and TVOR gains at both 2 and 8 months (Fig. S1), along with variable phase responses and substantial signal distortion. Thus, we find a progressive and durable normalization of vestibular reflexes following LY411575 treatment, with functional performance ultimately matching wild-type levels.

### Restoration of afferent innervation following LY411575 induced hair cell regeneration

To assess neural activity of vestibular afferents at regenerated hair cells, we measured spontaneous discharges and dynamic responses to head rotation and translation by single unit recording (Fig. 3A). A total of 498 afferents were recorded from the vestibular nerve bundles 8 months after treatment across three experimental groups: wild-type (n=190), DT-ablated without LY411575 (n=177) and DT-ablated with LY411575 (n=131). Afferents were classified, based on their responsiveness to head rotation and translation, as canal or otolith, and as regular (CV* < 0.1) or irregular (CV* > 0.1), based on the normalized coefficient (CV*) of variation of the interspike intervals (Fig. 3B).

**Fig. 3.**
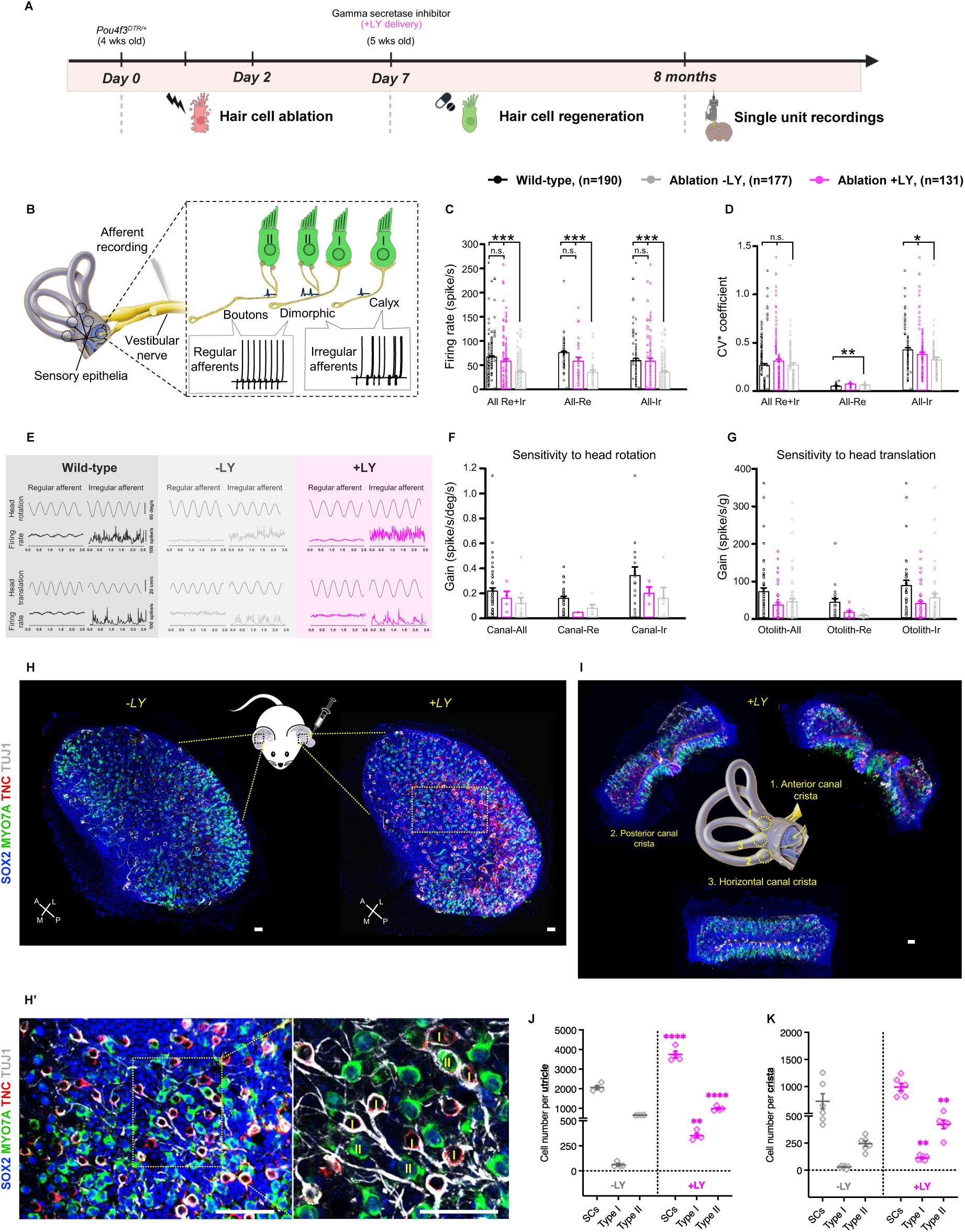
Sustained restoration of canal and otolith afferents at 8 months by γ-secretase inhibitor treatment. **(A)** Experimental design for drug delivery in DT-pretreated *Pou4f3^DTR/+^* mice. Single unit recording of vestibular afferents was performed 8 months after drug treatment +/-LY. **(B)** Drawing of vestibular afferents depicting a bouton ending of a regular afferent contacting a type II hair cell (left cell), a calyx ending of an irregular afferent around a type I hair cell (right cell), and a dimorphic afferent contacting both hair cell types (middle cells). Inserts show the difference in the resting discharge variability of regular and irregular afferents. **(C)** Averaged spontaneous firing rates from wild-type (black), *Pou4f3^DTR/+^* DT-ablated -LY (gray) and +LY (pink) mice. **(D)** Normalized coefficient of variation (CV*) of inter-spike intervals from the same groups showing the regularity of vestibular afferents. **(E)** Regular and irregular canal and otolith afferent responses to nasal-occipital head rotation and translation in wild-type (black), *Pou4f3^DTR/+^* DT-ablated -LY (gray) and +LY (pink) mice. **(F)** Canal afferents sensitivity to head rotation gains. **(G)** Otolith afferents sensitivity to head translation gains. **(H-I)** Sox2 (blue), Myo7a (green), Tnc (red) and Tuj1 (gray) immunolabelled sensory epithelia showing utricles from untreated (-LY) and treated (+LY) ears **(H)** with high-magnification images indicating afferents innervating type I (Myo7a+Tnc+) and type II (Myo7a+Sox2+) hair cells **(H’)** and cristae from drug treated ear **(I)**. **(J** *and* **K)** Quantification of supporting cells (SCs), type I and type II hair cells from untreated (-LY) and treated (+LY) utricles **(J)** and cristae **(K)**. All data represent the mean ± SEM. \*\*\*\**p* < 0.0001, *\*\*\*p* < 0.001, \*\**p* < 0.01 and \**p* < 0.05 by 1-way ANOVA with Tukey’s multiple comparisons test. Scale bars = 50-100 µm.

LY411575-treated mice exhibited substantial recovery of vestibular afferent function. Spontaneous firing rates were significantly elevated in both regular and irregular afferents compared to untreated DT-ablated animals, approaching wild-type levels (Fig. 3C). Firing regularity, assessed by CV* was similar in LY-treated and wild-type mice, indicating stable spike timing after regeneration of hair cells in the canal and otolith organs (Fig. 3D and E). Sensitivity to head motion was improved, with canal and otolith afferents in LY-treated mice showing clear responses to head rotation and translation (Fig. 3E-G). These neural response patterns aligned with the robust recovery of RVOR and TVOR observed following γ-secretase inhibition.

To further characterize the restoration of the vestibular circuits, we examined afferent morphology and hair cell reinnervation. The vestibular system comprises distinct nerve endings: i) regular afferents with bouton endings contacting type II hair cells, which encode fine motion cues at low frequencies; ii) irregular afferents with calyx endings surrounding type I hair cells, which convey rapid head motion at high frequencies; and iii) dimorphic afferents contacting both hair cell types (Fig. 3B). Immunolabeling of sensory epithelia, utricles (Fig. 3H) and cristae (Fig. 3I), using markers for Sox2, Myo7a, Tnc, and Tuj1 revealed reinnervation after LY411575 treatment. High-magnification images demonstrated afferent endings forming calyces and boutons with type I (Myo7a+Tnc+) and type II (Myo7a+Sox2+) hair cells, respectively (Fig. 3H’). Quantification of supporting cells and hair cells showed significantly higher cell numbers following LY411575 treatment (Fig. 3J and K). Drug-regenerated organs had an approximately 2-fold increase in total hair cells compared to untreated controls, with type I hair cells comprising 15-20% of the regenerated population. These results demonstrate that γ-secretase inhibition induces a comprehensive restoration of vestibular sensory function, encompassing cellular regeneration, neural circuit reconstruction, and functional recovery.

### Vestibular hair cell regeneration induced by Notch1 knockout

Treatment with a γ-secretase inhibitor was chosen as the therapeutic approach because of the known effect of inhibiting this enzyme on signaling pathways important for maintaining hair cell identity (*36, 40, 42*). However, there are more than 90 known γ-secretase substrates (*45*), and we sought to identify the substrate underlying these effects on the vestibular system.

To investigate whether *Notch1* was the target for the effect of the γ-secretase inhibitor on vestibular hair cell regeneration, we deleted *Notch1* in supporting cells of the *Pou4f3^DTR/+^* mouse crossed to a *Plp1^CreER^*;*tdTomato* mouse 2 days prior to DT-mediated ablation of hair cells. As *Plp1* expression is restricted to supporting cells in the utricular sensory epithelium (*46*), our approach allowed lineage tracing of Plp1⁺ supporting cells (Fig. 4A and B).

**Fig. 4.**
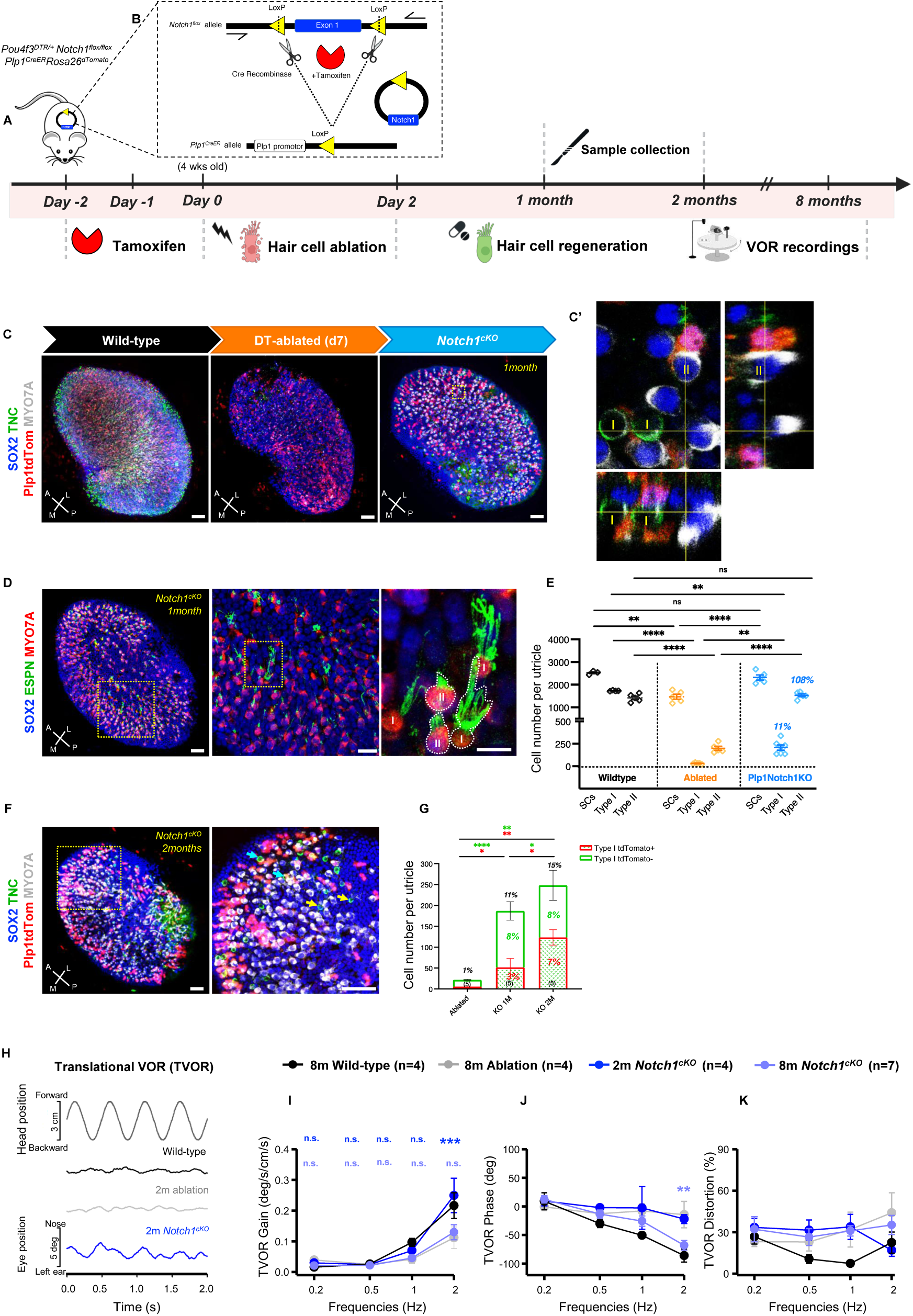
Conditional knockout of *Notch1* in Plp1+ supporting cells increases type I and II hair cell regeneration and restores VOR responses. (A) Schematic of the experimental approach, *Pou4f3^DTR/+^ Plp1^CreER^ Notch1^fl/fl^ tdTom* and wild-type were subjected to tamoxifen to induce Cre-mediated recombination in *Plp1^CreER^ floxed-Notch1* followed by diphtheria toxin (DT) to ablate hair cells. (**B)** The genetic strategy using Cre recombinase and tamoxifen to induce *Notch1* deletion, with LoxP sites flanking exon 1 of the *Notch1* gene. (C *and* C’) Utricles immunolabeled for Sox2 (blue), Tnc (green), Myo7a (green) with tdTomato (red) for fate mapping of supporting cells, high magnification images (**C’)** show an orthogonal view of newly regenerated type I (Myo7a+Tnc+) and type II (Myo7a+Tnc-Sox2+) hair cells arising from Plp1tdTom+ supporting cells. (**D)** 1-month-old *Pou4f3^DTR/+^ Notch1^cKO^* DT-ablated utricle immunolabeled for Sox2 (blue), Espn (green), Myo7a (red), high magnification images indicate the presence of stereociliary bundles in type I and type II hair cells. (**E)** Quantification of type I and type II hair cells and supporting cells. (**F)** 2-month-old *Pou4f3^DTR/+^ Notch1^cKO^* DT-ablated utricle immunolabeled for Sox2 (blue), Tnc (green), Myo7a (gray) with tdTomato (red) in Plp1 supporting cells. High magnification image indicates type I cells arising from Plp1tdTom+ (cyan arrows) and Plp1tdTom-(yellow arrows). (**G)** Type I hair cell numbers from DT-ablated *Pou4f3^DTR/+^*compared to *Pou4f3^DTR/+^ Notch1^cKO^ tdTom* mice 1 and 2 months after ablation. **(H)** Translational vestibuloocular reflexes (TVOR) in response to sinusoidal head translation at 2 Hz 2 months after ablation in wild-type (black), *Pou4f3^DTR/+^* DT-ablated (gray) and *Pou4f3^DTR/+^ Notch1^cKO^* DT-ablated (blue) mice. TVOR gains **(I)** phases **(J)** and distortions **(K)**. All data represent the mean ± SEM. *\*\*\*\*p* < 0.0001, \*\**p* < 0.01, \**p* < 0.05 by 2-way ANOVA with Tukey’s multiple comparisons test. Scale bars = 50-100 µm. See also Fig. S2 and Table S2.

At 1 month, *Notch1^cKO^* mice showed robust regeneration of Myo7a+ hair cells in the utricle. Immunolabeling revealed that new hair cells in the lineage-traced mice differentiated into both type I and type II hair cells (Fig. 4C). Tnc staining confirmed the presence of calyces surrounding the base of the type I cells, which were further identified through orthogonal views capturing the lineage tag in both type I and type II cells (Fig. 4C’). Espin staining highlighted distinct stereociliary morphologies (Fig. 4D), short and thin stereocilia characteristic of type II hair cells, and longer, thicker stereocilia characteristic of type I hair cells (*7, 29, 47*). Quantification indicated increased numbers of both hair cell types and supporting cells in *Notch1^cKO^*ears compared to ablated controls (Fig. 4E).

Two months after ablation, *Notch1* knockout resulted in a substantial increase in type I hair cells compared to 1 month (Fig. 4F and G). The regeneration was primarily observed in *Plp1tdTom+* cells, which increased from 3% of the type I cells at 1 month to 7% at 2 months, while the proportion of *Plp1tdTom*-type I cells remained stable at 8% (Fig 2G). As the *Plp1* promoter is inactive in supporting cells within the striolar region (*12*), a subset of type I cells must have been derived from a population of supporting cells that did not express *Plp1*. The presence of Sox2+ type I hair cells further supports a contribution from a distinct Sox2+ supporting cell population (Fig. 4C’).

The increased hair cell regeneration was associated with an improvement in the VOR responses (Fig. 4H and Fig. S2). At 2 months, *Notch1^cKO^* mice exhibited compensatory TVOR gains comparable to wild-type levels (Fig. 4I). TVOR phase (Fig. 4J) and distortion (Fig. 4K) measurements showed improved synchronization between head motion and eye movement. Longitudinal analysis revealed a decline in vestibular function between 2 and 8 months, with TVOR responses exhibiting reduced gain, particularly at higher frequencies *(****p= 0.0003,* 2Hz) (Fig. 4I and Table S2). RVOR measurements indicated partial functional recovery at 2 months, with gains (Fig. S2) at 8 months reaching wild-type levels in some cases (Table S2). Afferent recordings confirmed an increased number of rotational and translation afferents in *Notch1^cKO^*mice at 8 months, with spontaneous firing rates and CV* values comparable to wild-type mice (Fig. S3).

### Divergent long-term effects of pharmacological inhibition and genetic knockout of *Notch1*

Statistical analysis revealed important differences in vestibular recovery dynamics between LY411575-treated and *Notch1^cKO^* mice (Fig. S3). There were no significant differences in TVOR gain between the 2 approaches at 2 months, supporting comparable initial recovery (Fig. S3B). However, by 8 months, a significant decline in TVOR performance was observed in *Notch1^cKO^* at 2 Hz *(**p= 0.0077)*, with LY411575-treated mice showing superior recovery comparable to wild-type mice (Fig. S3B’). For RVOR gains, LY411575-treated mice had significantly improved performance compared to *Notch1^cKO^* mice at 2 months across high frequencies *(*p = 0.045, **p = 0.0048, **p = 0.001* and *\*\*\*\*p < 0.0001* for 0.5, 1, 2 and 4 Hz, respectively) (Fig. S3A and C). By 8 months, these differences were largely diminished, with statistical significance only remaining at 4 Hz *(*p = 0.0425)* (Fig. S3A’). These findings suggest that while both approaches enable hair cell regeneration, γ-secretase inhibition results in more stable long-term recovery, potentially related to differential effects on type I vs type II hair cell regeneration and connectivity.

## Discussion

A lack of treatments to achieve reversal is a persistent problem for balance disorders (*2*). We demonstrate here that inhibition of γ-secretase one week after ablation of hair cells results in the regeneration of hair cells and restoration of vestibular function across test frequencies. The γ-secretase inhibitor treatment not only induces hair cell regeneration but also successfully reestablishes functional neural circuits accurately encoding vestibular stimuli. The single-unit recordings provide critical electrophysiological evidence for the restoration of vestibular afferent signaling and directly corroborate the morphological and functional recoveries. Specifically, the enhanced spontaneous firing rates, preserved neural signal regularity, and recovered sensitivity to head movements align precisely with the regeneration of type I and type II hair cells.

Our results represent a clear advantage over previously achieved regeneration. Previous work in the adult has shown limited hair cell regeneration or functional restoration either spontaneously or by gene therapy (*15, 19, 27, 31, 48*). The regeneration of both type I and type II hair cells and their reinnervation by the type-specific vestibular afferents suggests a promising therapeutic approach for vestibular disorders.

The genetic deletion of *Notch1* also resulted in hair cell regeneration and functional improvement, indicating that *Notch1* was likely the target accounting for the effect of the γ-secretase inhibitor, an important point because of the large number of possible γ-secretase substrates (*45*). Notch signaling is a key regulator of hair cell differentiation and controls the proportion of hair cells and supporting cells made during development of the sensory epithelium (*33, 35*). Notch-mediated lateral inhibition directs some progenitor cells to become hair cells, while others differentiate into supporting cells (*33–35*). Inhibition of Notch triggers hair cell regeneration from supporting cells in the newborn cochlea (*36, 38, 40*) and in organoids (*39*), but regeneration is limited in the adult mouse cochlea (*41–43*). We hypothesized that the inhibition of Notch might enhance regeneration in the adult vestibular organs, where hair cells regenerate spontaneously early in life (*13*), and chromatin is more permissive than in the cochlea (*44*).

Efforts to reprogram supporting cells to hair cells in adult inner ear using transcription factors have also resulted in a low extent of regeneration (*19, 44, 49*). We favored the pharmacological manipulation of the Notch pathway over transcription factor overexpression for reprogramming because it initiated a cascade of transcription factor expression that resembled normal development, including *Atoh1* and downstream transcription factors like *Gfi1* and *Pou4f3* (*36, 40, 42, 50, 51*). Notch signaling is upstream of *Sox2* activity in cochlear cells (*52*) and more broadly in neural development (*28, 36, 40, 52–55*), and both *Sox2* and *Notch1* can determine hair cell fate (*36, 50*). Sox2 controls the level of key transcription factors in hair cell development. Indeed, Sox2 can destabilize Atoh1 by stimulating expression of its cognate E3 ligase (*56*), and lowering Sox2 level preserves Atoh1 and can drive hair cell differentiation (*57*). *Sox2* knockout during early development increased type I hair cells in the vestibular system, while decreasing type II hair cells (*28*), and the effect of Notch on Sox2 is thus a likely mechanism for the increased ratio of type I to type II hair cells after Notch inhibition.

The regeneration of mouse vestibular hair cells was achieved in a damage model where 92% of hair cells were ablated. Inhibition of γ-secretase one week after ablation of hair cells resulted in 15-20% replacement of type I and complete replacement of type II hair cells in mouse vestibular organs. The experiments were performed at 4 weeks of age, when hair cells are mature and spontaneous regeneration is minimal. The limited regeneration in the absence of drug treatment is consistent with results in adult damage models (*14, 16*) and was restricted to type II hair cells (*15, 25, 27, 48*). VOR and single unit recording showed γ-secretase inhibitor treatment restored canal and otolith activity across frequencies, consistent with extensive regeneration of type I and type II vestibular hair cells. The continued improvement in these metrics of vestibular function 8 months after treatment with the γ-secretase inhibitor diverged from the *Notch1^cKO^*, suggesting that brief inhibition by locally administered drug provided a more stable effect than *Notch1* knockout. This could be explained by the permanent loss of Notch activity in vestibular supporting cells in the *Notch1^cKO^*, consistent with previous observations that Notch activity is required for the development and maintenance of vestibular organs (*33, 34, 46*). Failure of γ-secretase inhibitors for the treatment of Alzheimer’s disease was due to complications when using daily doses of inhibitor (*58*); a short duration of diminished Notch activity by administering drug into the inner ear thus proved more effective than deletion of the gene and avoided potential systemic toxicities. Local administration of γ-secretase inhibitors into the cochlea (*42*) to achieve transient inhibition has been shown to give an exposure of the inner ear of less than 48 hrs (*37*), an exposure that appears from this work to be sufficient for treatment of balance disorders with a single injection of drug. In a previous clinical trial of a γ-secretase inhibitor for sensorineural hearing loss (*59*), the recovery of hearing was modest, but the small intratympanic doses of drug had significant effects on understanding of speech in background noise.

Our demonstration of normal VORs, a clinically relevant metric for vestibular function, in the γ-secretase inhibitor-treated animals indicates that restoration of cells in this complex sensory epithelium can be achieved by a brief lowering of Notch activity using a drug injected locally in the ear. This result is highly significant for the development of regenerative therapeutics for the treatment of this common disorder in older adults (*2, 3*).

## Materials and Methods

### Mice

All mice were obtained from the Jackson Laboratory (Table S3), housed with open access to food and water. For hair cell loss, *Pou4f3^DTR^* mice were mated with C57BL/6 mice to obtain *Pou4f3^DTR/+^ and Pou4f3^+/+^*. Wild-type mice were used as controls. *Pou4f3^DTR/+^* heterozygous mice were used for hair cell ablation.

For *Notch1* conditional knockout experiments, *Notch1^flox/flox^*mice were mated with *Plp1^CreER^* to generate *Notch1^flox/flox^;Plp1^CreER^*. Littermates was crossed with *Pou4f3^DTR/+^* to get *Pou4f3^DTR/+^;Notch1^flox/flox^; Plp1^CreER^*. The homozygous mutants were defined as *Notch1* knockout. Littermates without Cre were used as controls for the spontaneous regeneration. We used *Pou4f3^DTR/+^;Notch1^flox/flox^; Plp1^CreER^* crossed *Rosa26^tdTomato^* (called *tdTomato*) mice for lineage tracing experiments. Male and female mice were used in all experiments. When mated, all progenies were on C57Bl/6J background. Littermates were genotyped with Jackson Laboratory protocols

### Sex as a biological variable

Mice of both genders were used in this study.

### Study approval

All mouse experiments were approved by the Institutional Animal Care and Use Committee of Massachusetts Eye and Ear and the University of Mississippi Medical Center.

### Diphtheria toxin administration

Diphtheria toxin (Sigma D0564) was reconstituted in distilled water at 1 mg/ml as a stock solution and stored at -20°C. A working solution was diluted in sterile PBS 1X at 10 µg/ml and freshly prepared prior to each injection. 4-week-old mice received 2 intramuscular injections of DT at 50 ng/g, spaced 2 days apart. Mice were hydrated with 0.4 ml of lactated Ringer’s solution injected subcutaneously at day 0, day 1, and day 2. Food was substituted by high caloric one and hydrogel was provided in the cage.

### Tamoxifen injections

For Cre activation, tamoxifen (Sigma T5648) was dissolved in corn oil at 25 mg/ml and administered by intraperitoneal injection to *Pou4f3^DTR/+^;Notch1^flox/flox^; Plp1^CreER^*;*Rosa26^tdTomato^* mice for 2 consecutive days prior to DT treatment (0.125 µg/g of body weight).

### Round window administration of LY411575

5-week-old DT-ablated mice of both genders received a unilateral injection of γ-secretase inhibitor (LY411575) onto the round window niche (RW), the contralateral ear was used as a control for the spontaneous regeneration. The drug was diluted in polyethylene glycol 400 (1:2) and artificial perilymph (1:2) to a final concentration of 4 mM. Animals were anesthetized with an intraperitoneal injection of ketamine (100 mg/kg, ip) and xylazine (10 mg/kg, ip). All the surgical procedures were performed in pre-warmed room (26°C), to maintain the body temperature. The fur behind the left ear was shaved with a razor and sterilized with 10% povidone iodine. Under an operating microscope, an incision was made posterior to the pinna near the external meatus. The otic bulla was opened to approach the round window niche. LY411575 solution was injected with a Hamilton syringe and administrated for 2 min. Gelatin was placed on the niche to maintain the solution, and the wound was closed. The solution was administered for 2 min. This approach is widely used clinically and has the advantage of sparing the inner ear but still taking advantage of the local route provided by the round window membrane for delivery of drug into the inner ear (*60*). Hydrogel and food gel were placed into the cage daily from the first day after surgery until recovery. Pain was controlled with buprenorphine (0.05 mg/kg) given directly postoperatively and meloxicam (2 mg/kg) injected every 20-24 hours for 3 days. Recovery was closely monitored daily for at least 5 days postoperatively.

### Vestibulo-ocular reflex

Detailed methods were previously described (*61–63*). Briefly, each mouse was implanted with a head holder on the skull. Eye position signals of the left eye were recorded using an eye tracking system (ISCAN ETS-200, ISCAN, Burlington, MA) that was mounted on a servo-controlled rotator/sled (Neurokinetic, Pittsburgh, PA). The eye tracker tracked the pupil center and a reference corneal reflection at a speed of 240 frames per second with a spatial resolution of 0.1 deg. Calibration was achieved by rotating the camera from the left 10 degree to the right 10 degree around the vertical axis of the eye. Following calibration, horizontal head rotations were delivered at 0.2, 0.5, 1, 2 and 4 Hz (60 degree/s peak velocity) to measure the steady state rotational VOR responses. Signals related to eye position and head position were sampled at 1 kHz at 16 bits resolution by a CED Power 1401 system (Cambridge Electronics Devices, Cambridge, UK). Eye movement responses were analyzed using Spike2 (Cambridge Electronics Devices), MatLab (MathWorks, Natick, MA) and SigmaPlot (Systat Software, San Jose, CA). Eye position signals were filtered and differentiated with a band-pass of DC to 50 Hz to obtain eye velocity. Gains and phases of the rotational VORs were calculated by performing a fast Fourier transform on the de-saccaded eye velocity signal and head rotation velocity signal as described (*63*).

### Vestibular afferent recording

Single unit recording of vestibular afferents was performed under ketamine/xylazine anesthesia as described (*63–65*). The head was stabilized on a stereotaxic frame (David Kopf Instruments, Tujunga, CA, USA) via the head holder. The animals’ core body temperature was monitored and maintained at 36-37°C with a heating pad (Frederick Haer & Company, Bowdoinham, ME, USA). A craniotomy was performed to allow access of the vestibular nerve by a microelectrode filled with 3 M NaCl (40∼60 MΩ) (Sutter Instruments, Novato, CA, USA). Extracellular recording was performed using a MNAP system (Plexon Inc., Dallas, TX, USA). Every spontaneously active nerve fiber encountered was tested. Each afferent’s spontaneous activity was first recorded to calculate the regularity and baseline firing rate. Each semicircular canal was then brought into the plane of earth-horizontal rotation, and the isolated afferent’s response to head rotation along with horizontal and vertical head position signals were recorded. Extracellular voltage signals were sampled by a CED at 20 kHz with 16-bit resolution and a temporal resolution of 0.01ms. Head position signals were sampled at 1 kHz. Regularity of vestibular afferents was determined by calculating their normalized coefficient of variation of inter-spike intervals, i.e., CV*s. Vestibular afferents were classified as regular (CV*<=0.1) or irregular (CV*>0.1) units based on their CV* (*9, 66, 67*). To quantify an afferent’s responses to head rotation, the fundamental response was extracted from the averaged data using a fast Fourier transform analysis. Gains and phases relative to head velocity were calculated at 1 Hz. Given that many afferents in damaged mice showed minimal modulation during head movement, traditional gain metrics were insufficient for assessing signal significance in their discharge activities. To address this, we employed a distortion metric, defined as 1-(amplitude of the fundamental response)/(square root of the sum of the squared amplitudes of the first 10 harmonics), for a more statistically confident evaluation of head movement signals. This distortion metric, akin to “stimulus-response coherence”, is particularly effective for assessing vestibular afferent responses to head rotation and translation in animal models with damaged vestibular end organs resulting from genetic mutations, trauma and ototoxicity (*66*). An afferent with translation distortion <=30% is classified as a translation/otolith afferent. An afferent with rotation distortion <=30% and translation distortion>30% is classified as a rotation/canal afferent. An afferent with rotation distortion>30% and translation distortion>30% is classified as a no response afferent.

### Immunohistochemistry

Utricles were harvested and fixed for 30 minutes in 4% paraformaldehyde in PBS, pH 7.4 (Electron Microscopy Services) at room temperature. Unspecific binding was blocked with 10% donkey serum, 0.25% Triton X-100 in PBS for 1 hour at RT. Tissues were incubated overnight at 4°C with specific primary antibodies diluted in blocking solution (Table S3). Then utricles were rinsed (3X 5 minutes each) in PBS and incubated with the corresponding AlexaFluor secondary antibodies (2 hours at RT) (Table S3), nuclei were counterstained with DAPI (1:1000). Samples were mounted using Prolong Gold antifading (Life Technologies) on glass slides. Positive and negative controls were performed for all immunostaining. The images were acquired with Leica SP8 confocal microscopy and analyzed with Fiji-ImageJ.

### Cellular quantification

Cells were counted manually using graphic tools; Cell counter Plugins; of Fiji-ImageJ software. Total hair cell and supporting cell counts per utricle were performed from 20x z-stack images of 29790 μm^2^ using a grids method or from the whole sensory epithelium for the newly regenerated hair cells. The count from 29790 μm^2^ represents the quantification in fifteen grids (1986 μm^2^ each) randomly chosen from both striolar and extrastriolar regions that fell within the sensory epithelium. To get the total cell number per utricle, we computed the area of the sensory epithelium using ImageJ-Measure. We then multiplied the total area by the summed counts from fifteen grids and divided the result by the counted area (29790). Only cells that had a healthy-appearing DAPI-labeled nucleus located in the sensory epithelium were counted as positive.

### Statistical analysis

All statistical analyses were performed using GraphPad Prism Software Version 10 for MacOS (www.graphpad.com). Data are reported as mean ± SEM. 1-way ANOVA or 2-way ANOVA with Tukey post-test for the pairwise comparison. P-values were adjusted for multiple companions. Mixed-effects model was used for comparison across multiple measurements. Statistical significance was defined as *P < 0.05*.

## Acknowledgments

National Institutes of Health grant R01DC020322 (AE)

## Author contributions

Conceptualization: AE, HL

Methodology: AE, HL, HZ, WZ

Investigation: AE, HL, HZ, WZ

Visualization: AE, HL, HZ, WZ

Funding acquisition: AE

Project administration: AE

Supervision: AE, HL

Writing – original draft: AE, HL

Writing – review & editing: AE, HL, HW, WZ

---

**Fig. S1.**
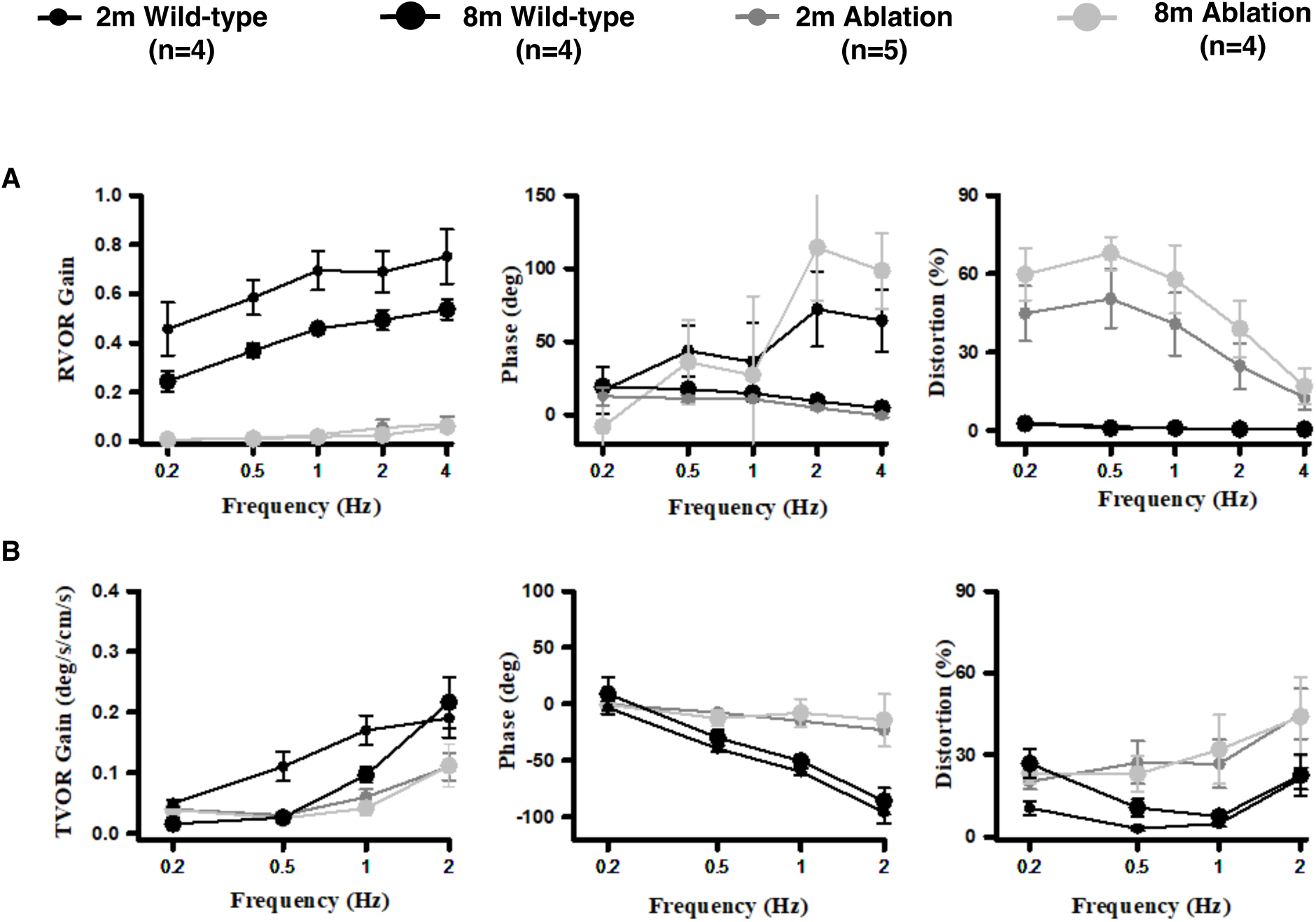
Vestibular reflex function is maintained over time in wild-type and spontaneous regeneration models. **(A)** Rotational vestibulo-ocular reflexes (RVOR) in response to head rotation, gains, phases and distortions. (**B)** Translational vestibulo-ocular reflexes (TVOR) in response to head translation, gains, phases and distortions. All data represent the mean ± SEM. VOR responses were comparable between 2 and 8 months for wild-type and spontaneous regeneration (DT-ablated untreated mice). Related to Figures 2 and 4.

**Fig. S2.**
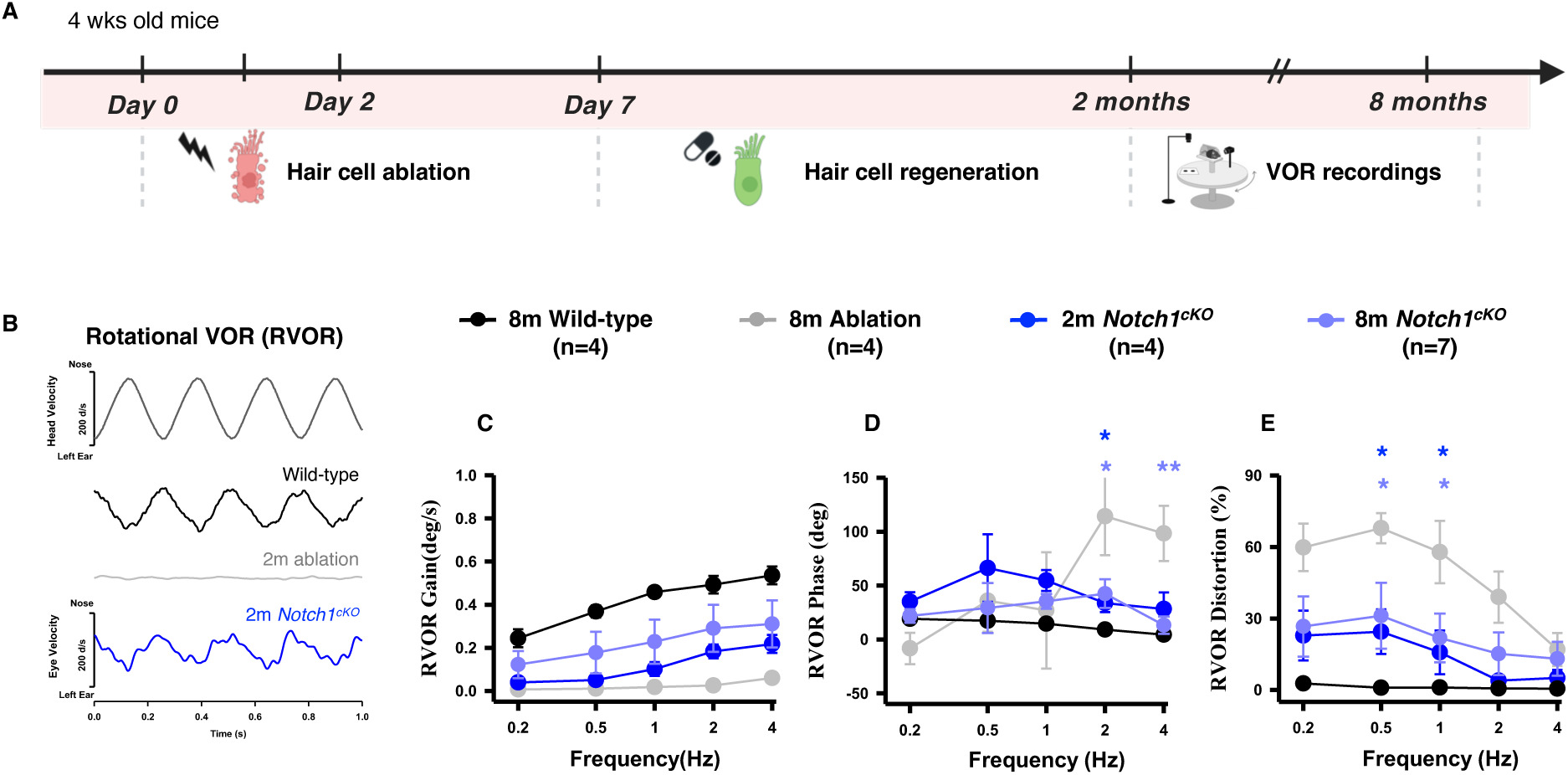
Improvement of rotational VOR responses in *Notch1^cKO^* DT-ablated mice. **(A)** Schematic of the experimental approach, Rotational VORs were recorded at 2- and 8-months post DT ablation. (**B)** Rotational vestibuloocular reflexes (RVOR) to sinusoidal head rotations with representative eye velocity responses to 4 Hz head rotation at 2 months, from wild-type (black), *Pou4f3^DTR/+^* DT-ablated (gray) and *Pou4f3^DTR/+^ Notch1^cKO^* DT-ablated (blue). RVOR gains **(C)** phases **(D)** and distortions **(E).** All data represent the mean ± SEM. \*\**p* < 0.01, \**p* < 0.05 by 2-way ANOVA with Tukey’s multiple comparisons test. Related to Figure 4.

**Fig. S3.**
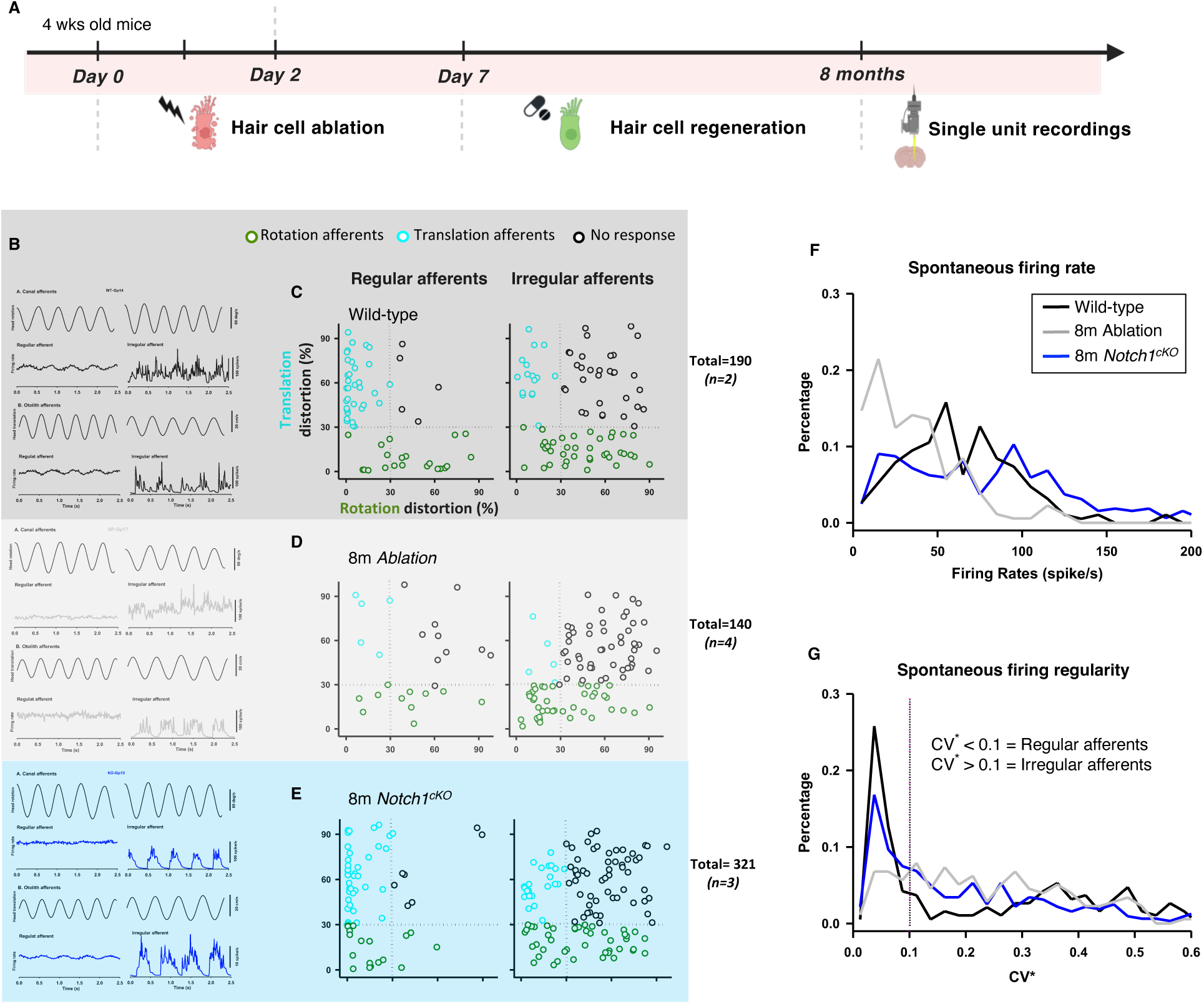
Restoration of vestibular afferent activity in *Notch1^cKO^* DT-ablated mice. **(A)** Schematic of the experimental timeline: single unit recording was performed at 8 months post-damage from 688 vestibular afferents in three groups: wild-type (190), DT-ablated (177), and *Notch1^cKO^*DT-ablated (321). (**B**) Regular and irregular canal and otolith afferent responses to nasal-occipital head rotation and translation from wild-type (black), *Pou4f3^DTR/+^* DT-ablated (gray) and *Pou4f3^DTR/+^ Notch1^cKO^* DT-ablated (blue). (**C-E**) Comparative analysis of regular and irregular afferent distribution in wild-type **(C)** DT-ablated **(D)** and *Notch1^cKO^*DT-ablated **(E)**, with x- and y-axes representing rotational and translational distortions respectively. (**F**) Spontaneous firing rates from the same groups. (**G**) Vestibular afferent regularity. Normalized coefficient of variation (CV*) of inter-spike intervals.

**Fig. S4.**
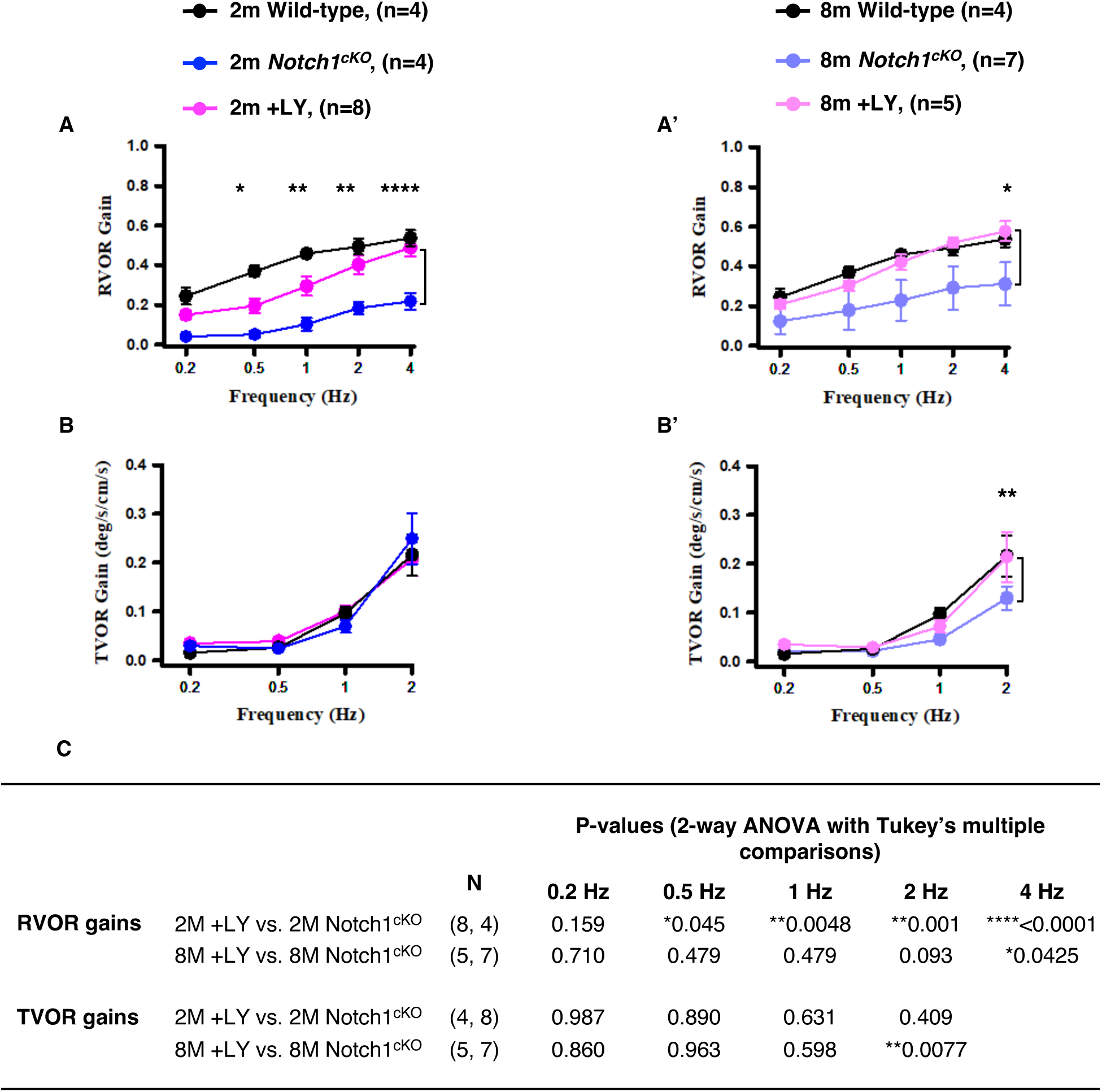
Comparison of VORs between the pharmacological and the conditional knockout approaches. **(A** and **B)** 2-month rotational (**A**) and translational (**B**) VOR gains. (**A’** and **B’)** 8-month rotational (**A’**) and translational (**B’**) VOR gains. (**C**) P-values for the comparisons between the pharmacological (+LY) and *Notch1* conditional knockout *(Notch1^cKO^)* approaches. All data represent the mean ± SEM. *\*\*\*\*p* < 0.0001, \*\**p* < 0.01, \*\**p* < 0.01 and \**p* < 0.05 by 2-way ANOVA with Tukey’s multiple comparisons; p-values reported in the graphics are from the comparisons of LY treatments to *Notch1^cKO^*.

**Table S1.** P-values for the comparisons between VOR gains for the DT-ablated without treatment (-LY), DT-ablated and drug treated (+LY) and wild-type animals. Related to. **Fig. 2**.

|  |  |  | P-values (2-way ANOVA with Tukey's multiple comparisons) |  |  |  |  |
| --- | --- | --- | --- | --- | --- | --- | --- |
|  |  |  | 0.2 Hz | 0.5 Hz | 1 Hz | 2 Hz | 4 Hz |
| RVOR gains | Wild-type vs. -LY | (4, 4) | ***0.0008 | ****<0.0001 | ****<0.0001 | ****<0.0001 | ****<0.0001 |
|  | Wild-type vs. 2M +LY | (4, 8) | 0.1909 | **0.0050 | **0.0086 | 0.2170 | 0.6951 |
|  | Wild-type vs. 8M +LY | (4, 5) | 0.8558 | 0.5342 | 0.8402 | 0.9430 | 0.8303 |
| TVOR gains | Wild-type vs. -LY | (4, 4) | 0.8440 | >0.9999 | 0.2889 | *0.0134 |  |
|  | Wild-type vs. 2M +LY | (4, 8) | 0.8599 | 0.9491 | 0.9989 | 0.9803 |  |
|  | Wild-type vs. 8M +LY | (4, 5) | 0.8881 | 0.9995 | 0.8010 | 0.9998 |  |

**Table S2.**
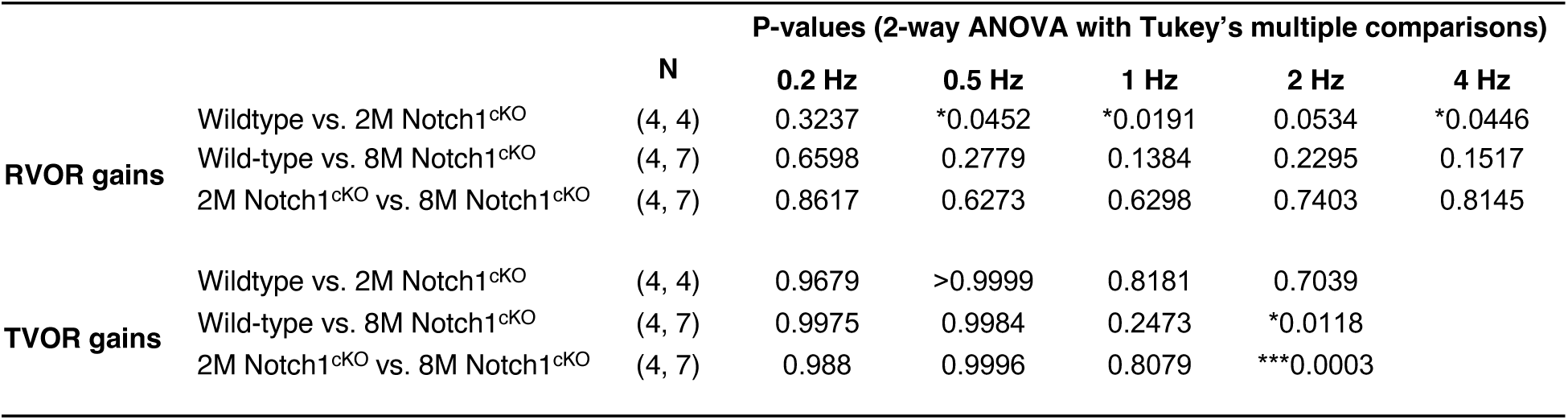
P-values for the comparisons between the VOR gains for the *Notch1*conditional knockout *and* the wild-type animals. Related to. **Fig.4**.

**Table S3:**
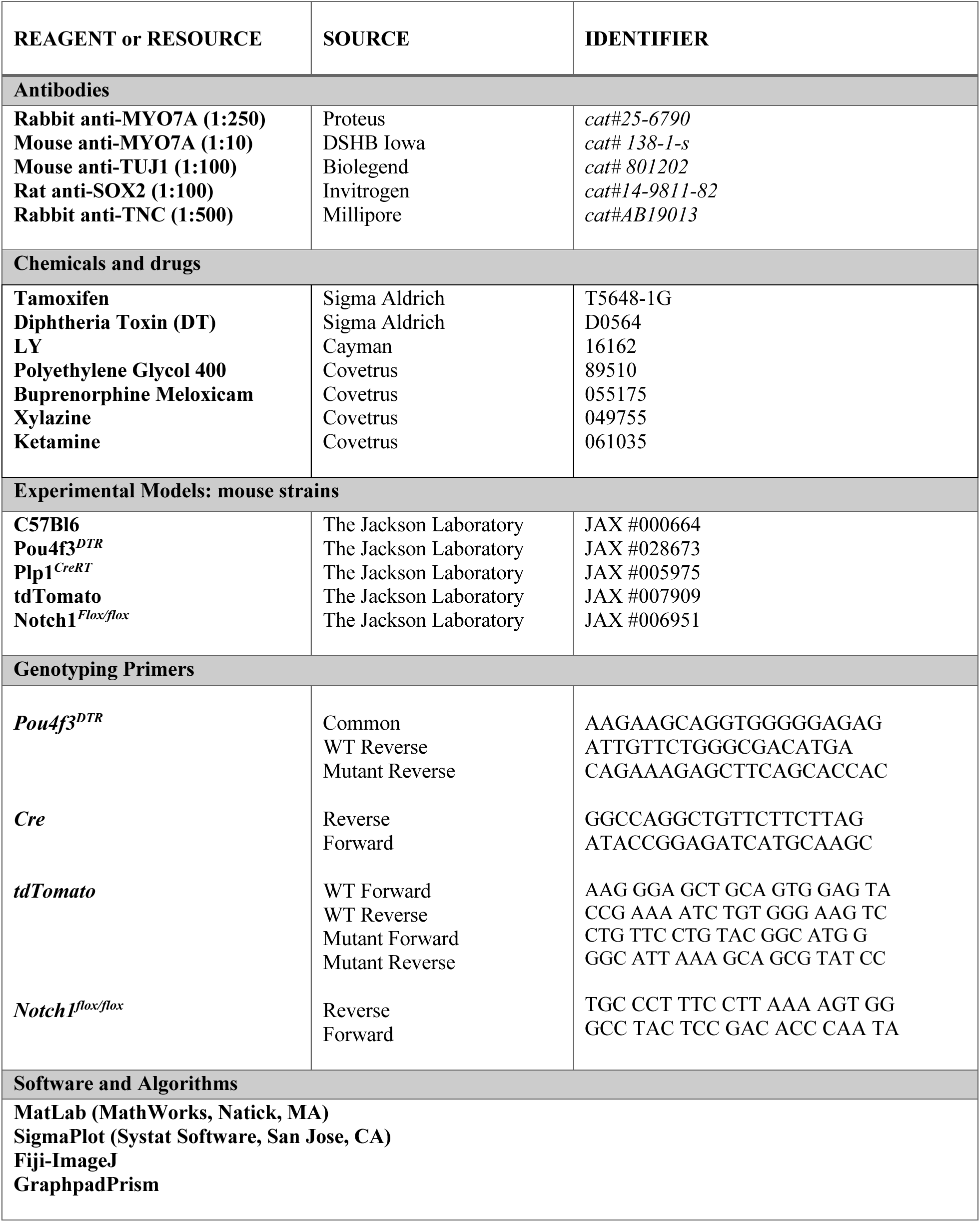
Resources.

